# Disruption of a *RAC1*-centred protein interaction network is associated with Alzheimer’s disease pathology and causes age-dependent neurodegeneration

**DOI:** 10.1101/713222

**Authors:** Masataka Kikuchi, Michiko Sekiya, Norikazu Hara, Akinori Miyashita, Ryozo Kuwano, Takeshi Ikeuchi, Koichi M. Iijima, Akihiro Nakaya

**Affiliations:** Department of Genome Informatics, Graduate School of Medicine, Osaka University, Osaka, Japan; Department of Alzheimer’s Disease Research, Center for Development of Advanced Medicine for Dementia, National Center for Geriatrics and Gerontology, Aichi, Japan; Department of Experimental Gerontology, Graduate School of Pharmaceutical Sciences, Nagoya City University, Aichi, Japan; Department of Molecular Genetics, Brain Research Institute, Niigata University, Niigata, Japan; Asahigawaso Research Institute, Asahigawaso Medical-Welfare Center, Okayama, Japan

**Author notes:** To whom correspondence should be addressed: (1) Masataka Kikuchi, 2-2 Yamadaoka, Suita, Osaka 565-0871, Japan, Email Address (MK), (2) Koichi M. Iijima, 7-430 Morioka-cho, Obu, Aichi 474-8511, Japan, Email Address (KMI). Equal contributors (Masataka Kikuchi, Michiko Sekiya, Norikazu Hara).

## Abstract

The molecular biological mechanisms of Alzheimer’s disease (AD) involve disease-associated cross-talk through many genes and include a loss of normal as well as a gain of abnormal interactions among genes. A protein domain network (PDN) is a collection of physical bindings that occur between protein domains, and the states of the PDNs in patients with AD are likely to be perturbed compared to those in normal healthy individuals. To identify PDN changes that cause neurodegeneration, we analysed the PDNs that occur among genes co-expressed in each of three brain regions at each stage of AD. Our analysis revealed that the PDNs collapsed with the progression of AD stage and identified five hub genes, including *Rac1*, as key players in PDN collapse. Using publicly available gene expression data, we confirmed that the mRNA expression level of the *RAC1* gene was downregulated in the entorhinal cortex (EC) of AD brains. To test the causality of these changes in neurodegeneration, we utilized *Drosophila* as a genetic model and found that modest knockdown of *Rac1* in neurons was sufficient to cause age-dependent behavioural deficits and neurodegeneration. Finally, we identified a microRNA, hsa-miR-101-3p, as a potential regulator of *RAC1* in AD brains. As the Braak neurofibrillary tangle (NFT) stage progressed, the expression levels of hsa-miR-101-3p were upregulated specifically in the EC. Furthermore, overexpression of hsa-miR-101-3p in the human neuronal cell line SH-SY5Y caused *RAC1* downregulation. These results highlight the utility of our integrated network approach for identifying causal changes leading to neurodegeneration in AD.

## Introduction

Alzheimer’s disease (AD) is a multifactorial disease caused by genetic and environmental factors, including ageing. AD is characterized by extracellular deposition of senile plaques (SPs) and intracellular accumulation of neurofibrillary tangles (NFTs). These aggregates sequentially spread across brain regions, and this development is described by the Braak SP or NFT stage (1). As the Braak NFT stage progresses, NFTs diffuse from regions such as the entorhinal cortex (EC) and hippocampus to the neocortical regions.

As brain pathologies progress, AD-associated molecular changes are induced in the brain, and these changes are thought to cause synaptic degeneration and neuronal death. To systematically detect such changes, genome-wide gene expression analyses have been carried out and have identified numerous genes differentially expressed in the AD brain, some of which are associated with the progression of SP or NFT pathology (2–7). These findings suggest that the progression of AD can be described as alterations in these molecular networks and that the elucidation of such changes will lead to the development of novel therapeutics for AD.

Network analysis is effective not only for detecting differences in molecular networks between AD patients and healthy controls but also for identifying hub genes that interact with many genes as key drivers of pathologies (8–14). A protein interaction network (PIN) is a collection of physical protein-protein interactions validated by high-throughput techniques, such as yeast two-hybrid systems and mass spectrometry-based technologies. PINs have diverse structures among different tissues and cell types and between normal and disease states (15,16). Such diversities may stem from differential gene expression patterns or differential exon usage by alternative splicing. Since the molecular mechanisms of AD involve a loss of normal as well as a gain of abnormal interactions among genes, we sought to systematically detect alterations in the physical associations of gene products associated with AD pathology.

In this study, we developed a network analysis method via a combination of PINs and gene expression data from AD brains. We constructed domain-level PINs (called protein domain networks (PDNs)) expressed in each Braak NFT stage by integrating the whole-exome expression data measured by exon array and domain-domain interaction data from the INstruct database (17). The INstruct database provides detailed domain-domain interaction data that combine protein physical interaction data and protein co-crystal structure data. Our analyses revealed that the loss of PDNs occurred with the progression of Braak NFT stage and identified *RAC1* as a hub gene, a key player in the alteration of PDNs. Using the fruit fly *Drosophila*, we experimentally validated that modest knockdown of *RAC1* in brain neurons induced age-dependent neurodegeneration. Our data further suggested that a microRNA (miRNA)-mediated process is involved in the downregulation of *RAC1* expression in response to the progression of Braak NFT stage.

## Results

In this study, we aimed to detect the dynamics of PDNs across Braak NFT stages in AD brains and to identify key networks and hub genes associated with the progression of Braak NFT stage (summarized in **Figure 1**). To this end, we re-analysed our exon array data from 71 postmortem brain samples, including three brain regions (the EC, temporal cortex (TC), and frontal cortex (FC)) at various Braak NFT stages (**Figure 1A**) (7). We then constructed PDNs in the three brain regions at each Braak NFT stage (**Figure 1B, left**). The structures of the PDNs were analysed to detect changes in the PDNs associated with the progression of Braak NFT stage (**Figure 1B, centre)**. The hub genes, which are connected to many genes, were identified in the PDNs in each brain regions (**Figure 1B, right**). Finally, we examined whether the identified hub genes were downregulated in independent mRNA expression data sets from AD brains (**Figure 1C, left**) and whether loss of function of the identified hub gene was sufficient to cause neurodegeneration (**Figure 1C, centre)**. We further identified miRNAs that could regulate the expression levels of the hub genes using bioinformatics tools followed by experimental validation using human neuronal cultured cells (**Figure 1C, right**).

**Figure 1.**
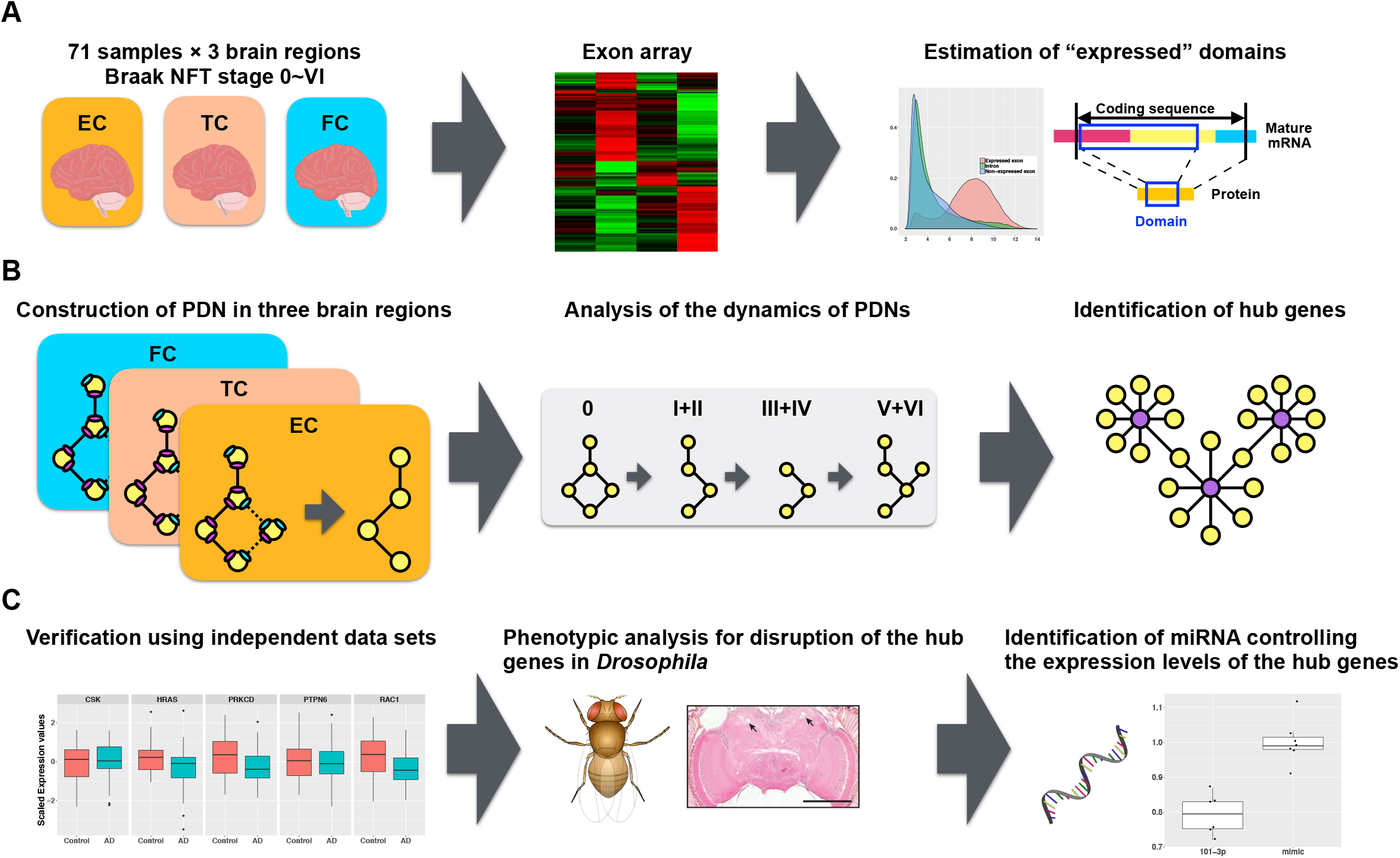
Overview of the present study. (A) Analysis of gene expression data to estimate “expressed” domains. Postmortem brain samples were obtained from three brain regions (the EC, TC, and FC) of 71 individuals. These samples were analysed by exon array. Expressed exons and domains in each brain region at each Braak NFT stage were estimated based on the expression values and statistics. (B) Network analysis to detect hub genes. Expressed domains were projected onto the domain-domain interaction data to construct the PDNs in each brain region at each Braak NFT stage. The structures of the PDNs were analysed to detect changes in PDNs associated with the progression of Braak NFT stage. Hub genes were identified from the PDNs to obtain AD-associated genes. (C) Validation of the hub genes. The expression changes in the hub genes were verified using independent data sets. The confirmed hub genes were characterized for age-dependent behavioural deficits and neurodegeneration using transgenic flies. Finally, miRNAs that regulate the expression levels of the hub genes were identified and validated in the human neuronal cell line SH-SY5Y.

### Construction of PDNs in the EC, TC, and FC at each Braak NFT stage

To compare the network structures across Braak NFT stages, we first constructed “expressed” PDNs in the three brain regions at each Braak NFT stage. As a reference for the potential PDNs in the brains, we utilized the INstruct database, which includes physical interactions between 3,626 human proteins based on co-crystal structures. These interaction data are composed of physical associations between protein domains (e.g., an interaction between domain A of protein A and domain B of protein B (**Figure 2A**)). To infer whether given protein interactions occur at each Braak NFT stage in each brain region, we used gene expression data from postmortem brain samples from patients with AD. We predicted that two proteins could potentially interact if the known binding domains of two genes were co-expressed in the same brain region at the same Braak NFT stage. Conversely, if one or both of the binding domains between two genes were not expressed, we predicted that these two proteins could not interact (**Figure 2B**). Exon array analysis was carried out in the postmortem brain samples obtained from the three brain regions of the 71 individuals. The samples were divided into four groups based on Braak NFT stage (stage 0 (n=13), stages I and II (n=20), stages III and IV (n=19), and stages V and VI (n=19)). An exon array is a DNA microarray with probesets for exons in the human genome and enables comprehensive quantification of changes in exon usage.

**Figure 2.**
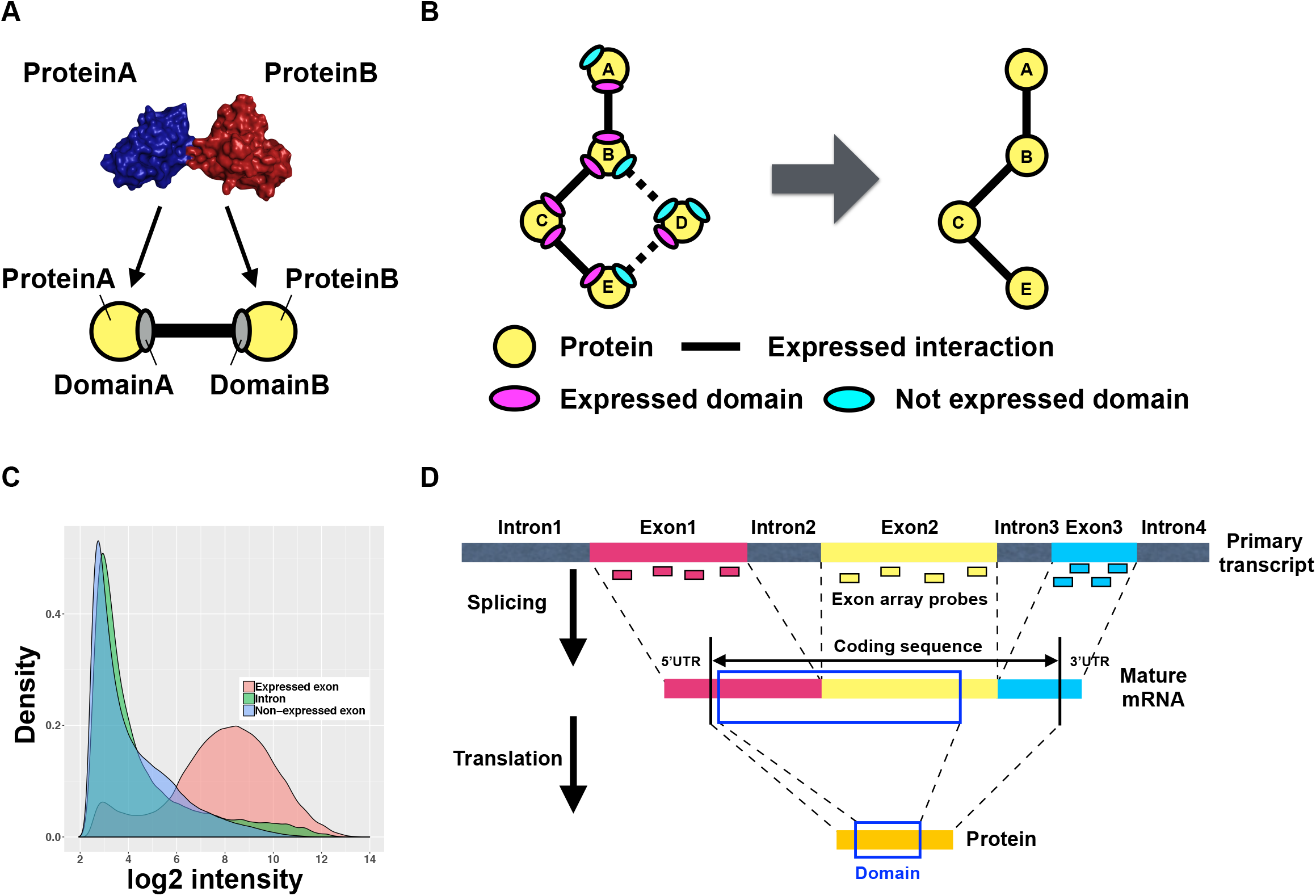
Construction of PDNs. (A) Schematic of a domain-domain interaction. The yellow circles indicate proteins. An ellipse indicates a domain. For instance, protein A and protein B interact through a physical association between domain A and domain B. (B) Identification of expressed PDNs. The ellipses in magenta and cyan indicate an expressed domain and a domain that is not expressed, respectively. If both domains that form an interaction are expressed, the interaction is assumed to be expressed. (C) Discrimination of expressed domains. A density plot shows the signals of all probes from expressed exons (red) and introns (green) and from non-expressed exons. The threshold determining whether a domain is expressed was set at 6. (D) Assignment of exons to a domain. The short rectangles indicate probes in each exon. The short rectangles outlined in blue show a domain in the corresponding protein sequence. In (D), the domain includes exon 1 and exon 2. Therefore, the expression level of the domain is determined by the expression levels of those exons.

We defined an “expressed” exon as an exon with a median detection above background (DABG) p-value of < 0.05 and a median log2-transformed expression value of ≥6 across all samples in a brain region at a Braak NFT stage. The DABG p-value is a metric used to detect signals that are distinct from background noise. The DABG p-value can highly discriminate between exons (assumed to be present) and introns (assumed to be absent) (18). A cutoff value of 6 was used based on a report by Kang *et al.* (19), which successfully explained gene expression dynamics across brain regions and ageing. Consistent with this report, the expression levels of probes in expressed exons were clearly distinct from those in non-expressed exons or introns in our data (**Figure 2C**). The area under the curve (AUC) was used as a measure of the ability of a metric to discriminate between the expressed exons and introns. The AUC for the above criteria (AUC=0.855) was higher than that for only the DABG p-value (AUC=0.691) (**Supplementary Figure S1**).

We examined exon usage and found that more than 90% of exons were stably expressed (i.e., satisfied the criteria described above) across all Braak NFT stages in all brain regions. This finding suggests that the majority of exons do not dynamically change with the progression of Braak NFT stage (**Supplementary Figure S2A**). In addition, more than 20% of genes had one or more domains that varied across Braak NFT stages (**Supplementary Figure S2B**). Each exon was assigned to the corresponding protein domain based on the DNA sequences and the amino acid sequences (**Figure 2D**) (see the Materials and Methods section for details). Finally, we identified the PDNs expressed in each Braak NFT stage by integrating the whole-exome expression data and the domain-domain interaction data from the INstruct database. As defined above, if all exons corresponding to a domain interaction between two proteins were “expressed”, we predicted that the interaction was “expressed”. In this way, we constructed a total of 12 PDNs expressed in each of the four Braak NFT stages in each of the three brain regions.

### PDNs altered with the progression of Braak NFT stage

To detect changes in PDNs associated with the progression of Braak NFT stage, we compared the expression profiles of PDNs in each brain region across Braak NFT stages. We focused on the subsets of PDNs whose expression patterns were positively (“appearing” PDNs) or negatively (“disappearing” PDNs) correlated with the progression of Braak NFT stage. These subsets of PDNs were classified into six expression patterns (**Figure 3A**). **Figure 3B and 3C** show the proportion of interactions constructing the appearing or disappearing PDNs against the total number of interactions in the PDNs in **Figure 3A**, respectively. We found that the proportions of disappearing PDNs were much higher than those of appearing PDNs in all brain regions, indicating that PDNs collapse with AD progression. Among the three patterns in disappearing PDNs, the most common pattern was “Pattern 1”, in which PDNs disappeared after Braak stage 0 (**Figure 3C**). In addition, the EC was the most severely affected among the three brain regions (**Figure 3C**). These results are consistent with the observation that the EC is affected more severely from the early stage of AD than the TC and FC (1).

**Figure 3.**
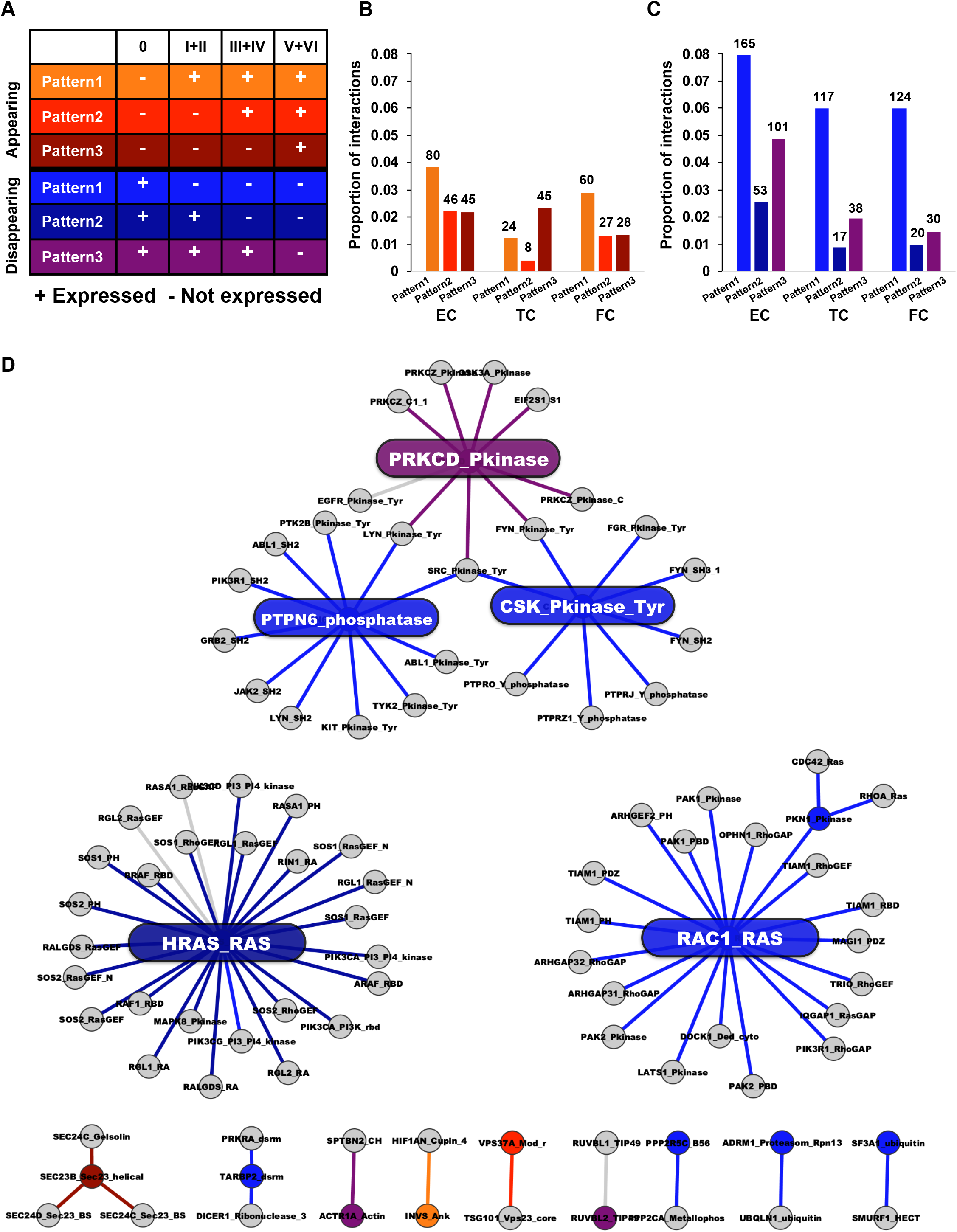
PDNs collapse with the progression of Braak stage. (A) Expression pattern of domain-domain interactions. “Appearing” and “disappearing” PDNs had three patterns each. Each pattern was defined by whether it was expressed or not in each Braak stage. (B, C) The proportions of interactions in appearing (B) and disappearing (C) PDNs. The colours in each bar correspond to the colours in the patterns in (A). The values on each bar indicate the number of interactions. (D) Braak stage-specific network perturbation in the EC. Each node and link indicate a protein domain and a domain-domain interaction, respectively. Those colours correspond to the expression patterns shown in (A). The labels on each node indicate the protein name and domain name. The coloured nodes and interactions were perturbed in a specific Braak stage but not by ageing. The grey colour indicates the other expression patterns.

To determine whether each pattern occurs by chance, we further examined the statistical significance relevant to the number of interactions constructing the PDNs in each expression pattern. We calculated theoretical values based on the probabilities of interactions expressed in each Braak NFT stage (see the Materials and Methods section for details). These theoretical values are equal to the proportion of interactions that occur by chance. We attempted to show whether the proportions of the observed interactions are greater or less than the theoretical values. “Pattern 1” in disappearing interactions was more than 44.6-fold higher than the theoretical value (48.4-fold in the EC, 44.6-fold in the TC, and 47.3-fold in the FC) (**Supplementary Table S1**). Next, we performed permutation tests with 10,000 replicates to calculate the p-values in each brain region (**Supplementary Figure S3**). The p-values of “Pattern 1” in disappearing interactions were significant across all brain regions (p-value < 1.00×10^−300^ in all brain regions).

We also found a statistically significant high number of interactions for “Pattern 3” in appearing interactions. “Pattern 3” is an expression pattern that was not present through Braak stages III+IV and was present only in Braak stages V+VI (17.2-fold and p-value = 2.70×10^−66^ in the EC; 23.4-fold and p-value = 9.08×10^−121^ in the TC; 9.34-fold and p-value = 9.56×10^−21^ in the FC). Comparison of the fold changes calculated above showed that “Pattern 1” in disappearing interactions had higher fold changes than “Pattern 3” in appearing interactions across all brain regions (2.81-fold (=48.4/17.2) in the EC, 1.91-fold (=44.6/23.4) in the TC and 5.06-fold (=47.3/9.34) in the FC). Taken together, our statistical results further confirmed the abovementioned collapse of PDNs.

To gain insight into the functional consequences of altered PDNs in each brain region, we performed gene functional enrichment analysis (**Supplementary Tables S2-7**). In the EC, the genes within appearing or disappearing PDNs were enriched in a pathway related to the immune systems (the top 11 in appearing PDNs (p-value = 5.43×10^−21^) and the top 10 in disappearing PDNs (p-value = 1.18×10^−38^)). In addition, the genes within disappearing domains were enriched in a phagocytosis-associated pathway (top 8 (p-value = 5.43×10^−21^)), suggesting impairment of phagocytosis in the EC. In the TC, the genes within appearing PDNs (top 39 (p-value = 4.89×10^−14^)) and those within disappearing PDNs (top 6 (p-value = 6.96×10^−28^)) were significantly enriched in a pathway related to axon guidance. We also found that the immunological pathway was enriched in the FC (the top 4 in appearing PDNs (p-value = 4.35×10^−29^) and the top 23 in disappearing PDNs (p-value = 7.37×10^−24^)).

### Identification of hub genes in PDNs that collapsed with the progression of Braak NFT stage in the EC

To identify key genes that play major roles in the perturbation of the PDNs, we searched for genes with hub domains, which have many interactions with other domains in each PDN. The patterns of those domains were classified into six expression patterns, similar to the classification shown in **Figure 3A**.

In general, Braak NFT stage correlates with ageing, suggesting that ageing is a potential confounding factor. Since we were interested in identifying AD-associated genes specifically related to Braak NFT stage rather than ageing, we extracted hub domains in which the expression patterns were specifically associated with the progression of Braak NFT stage but not with ageing using a linear regression model (those with an FDR-adjusted p-value for Braak NFT stage of < 0.05 and an FDR-adjusted p-value for ageing of > 0.05 in a Z-test). By these criteria, we identified 15 domains in only the EC (**Figure 3D**). Among the genes with these domains, we found five genes with hub domains (hereafter referred to as hub genes) having eight or more interactions. Interestingly, these hub genes were the disappearing genes.

*HRAS* (the HRas proto-oncogene, a GTPase) had the most disappearing interactions among the hub genes. *HRAS* belongs to the Ras family and is an oncogene. Upregulation of *HRAS* negatively controls the expression of the *CLU* gene, which was identified in some AD genome-wide association studies (GWASs) (20–22). *RAC1* is a member of the Rho family of small GTPases that regulates actin cytoskeleton remodelling and controls the formation of axon growth and guidance (23–25). The remaining three hub genes (*PTPN6, PRKCD*, and *CSK*) formed a network module. Protein Tyrosine Phosphatase, Non-Receptor Type 6 (*PTPN6*) is related to endocytosis along with *CD33*, a mutation in which was identified as a member of the susceptibility loci for AD by GWASs (26). Protein Kinase C Delta (*PRKCD*) is a PKC family member that has a role in learning and memory (27,28), and its activity is decreased in human AD brains (29,30). C-Terminal Src Kinase (*CSK*) acts as a non-receptor tyrosine protein kinase, and studies have suggested that *CSK* directly binds to N-Methyl-D-Aspartate receptor and controls synaptic plasticity (31,32).

Taken together, these reports suggest that the collapse of these networks may impair critical functions such as endocytosis and synaptic plasticity.

### A reduction in the mRNA expression levels of *RAC1* in AD brains was verified using independent data sets

To examine whether decreased expression levels of the hub genes identified in our data set could be verified with independent data sets, we used AlzData, a database providing statistical results from a meta-analysis using publicly available data sets, to re-analyse the expression levels of the five hub genes—*CSK, HRAS, PRKCD, PTPN6*, and *RAC1—*in the EC of patients with AD. Among the five genes, *RAC1* was significantly downregulated in the EC of AD patients (FDR-adjusted p-value = 0.046) (**Figure 4A**). Moreover, consistent with the results from our data sets, a reduction in the *RAC1* expression level was not simply correlated with ageing (**Figure 4B**). These results suggest that disruption of *RAC1* functions is specifically associated with AD pathology and may be a causative factor for neurodegeneration.

**Figure 4.**
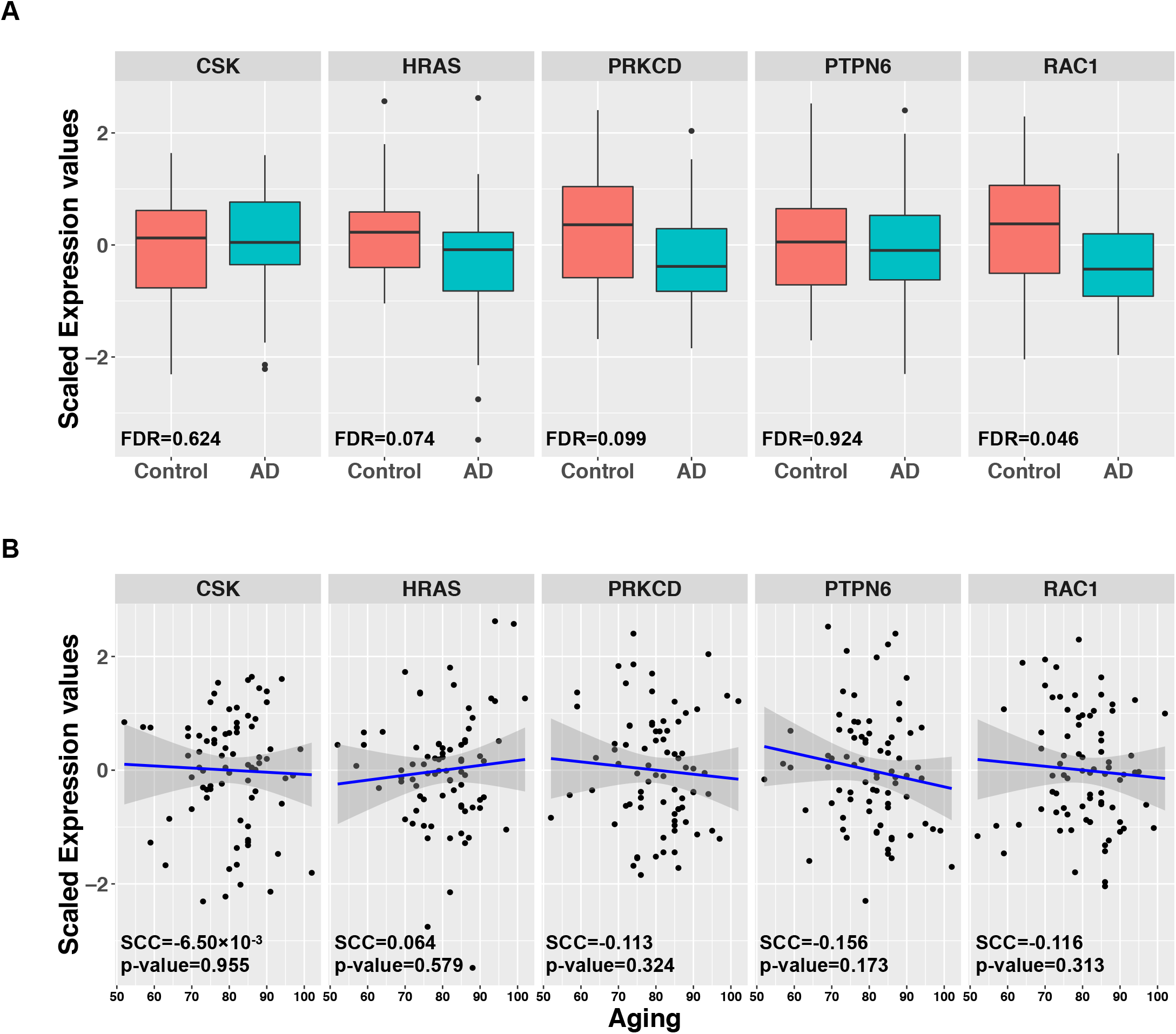
Expression levels of the five hub genes from a publicly available database, AlzData. (A) Comparisons of the expression levels of the five hub genes between control and AD in the EC region. FDRs were calculated based on p-values adjusted for age and sex by using a linear regression model. (B) Correlations between age and the expression levels of the five hub genes. SCCs were calculated. The p-values were calculated by a test for no correlation.

### Neuronal knockdown of *Rac1* causes age-dependent behavioural deficits and neurodegeneration

To determine whether a reduction in the RAC1 level is sufficient to cause age-dependent neurodegeneration, we utilized *Drosophila* as an *in vivo* genetic model. According to the *Drosophila* RNAi Screening Center (DRSC) Integrative Ortholog Prediction Tool (DIOPT), the primary amino acid sequences of *Drosophila Rac1* and human *RAC1*, which are similar in size (192 and 211 amino acids, respectively), exhibit 83% identity and 87% similarity.

To modestly knock down *Rac1* expression in fly neurons, the pan-neuronal elav-GAL4 driver was used to express shRNAi targeting *Rac1.* Quantitative RT-PCR analysis detected significant reductions (60%) in the mRNA expression level of *Rac1* in fly brains (Transgenic RNAi Project (TRiP) RNAi line #34910, **Figure 5A**). We first examined the effects of *Rac1* knockdown on locomotor functions in flies, as assessed by a forced climbing assay (33,34). Neuronal knockdown of *Rac1* significantly exacerbated age-dependent locomotor deficits after the age of 28 days (**Figure 5B**). In contrast, *Rac1* knockdown did not affect the life span of flies, suggesting that the observed locomotor deficits are not simply due to the reduced viability of flies (**Figure 5C**).

**Figure 5.**
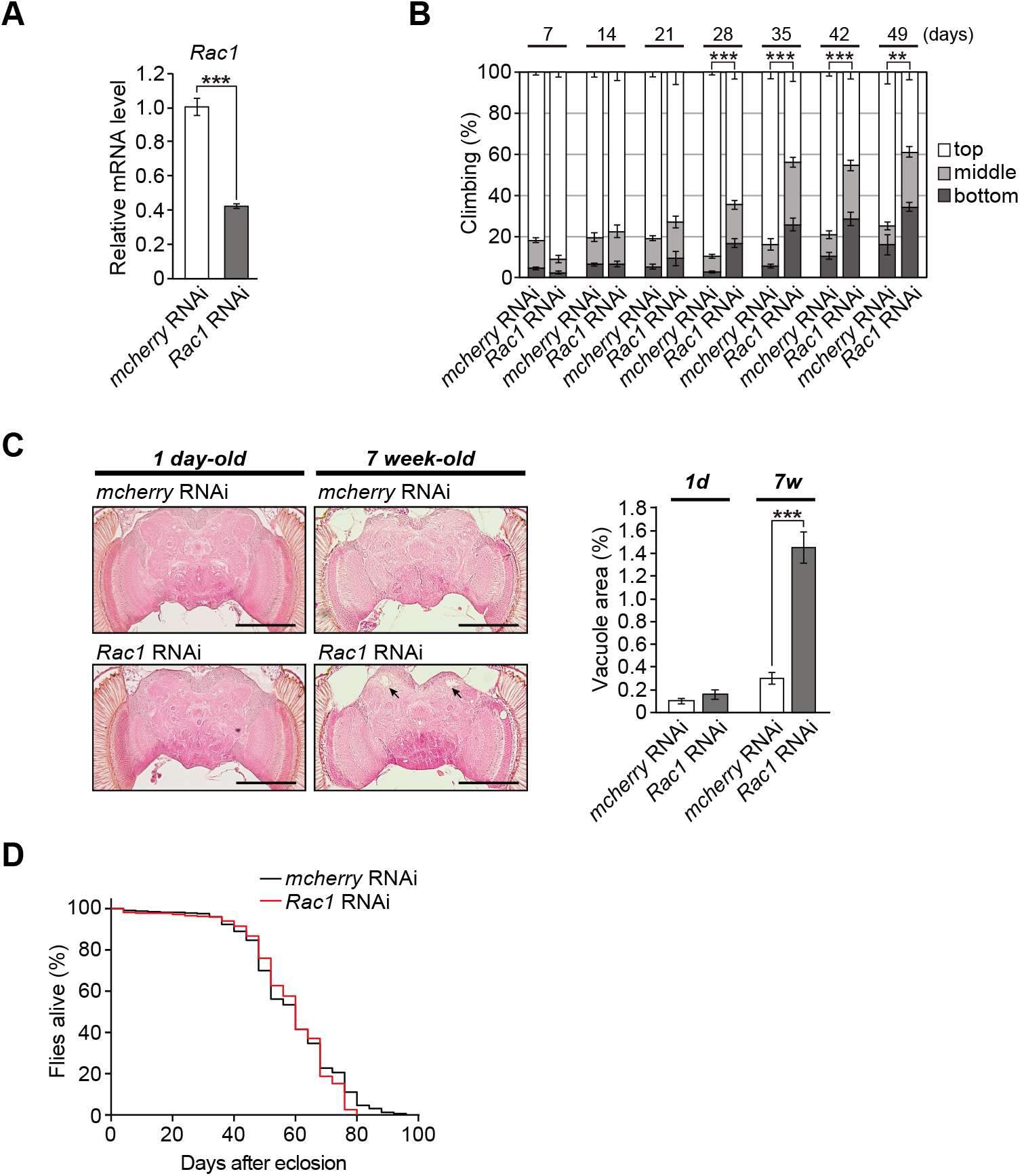
Neuronal knockdown of Rac1 causes age-dependent behavioural deficits and neurodegeneration. (A) mRNA levels of *Rac1* in the heads of flies carrying the RNAi transgene targeting *Rac1* were analysed by qRT-PCR. n = 4; ***p < 0.001 by Student’s *t*-test. (B) Knockdown of *Rac1* in neurons induced age-dependent locomotor deficits, as revealed by the climbing assay. The average percentages of flies that climbed to the top (white), climbed to the middle (light grey), or stayed at the bottom (dark grey) of the vials were calculated. The percentage of flies that stayed at the bottom was used for statistical analyses. n = 6 independent experiments; **p < 0.01 and ***p < 0.001 by Student’s t-test. (C) Knockdown of *Rac1* in neurons did not affect life span (n = 316, *Rac1* RNAi group; n = 326, control group). The life span of the flies was determined by Kaplan-Meier survival analysis with a log-rank test (n.s., not significant). (D) Knockdown of *Rac1* in neurons caused age-dependent neurodegeneration in the calyx (dendrite) region of the central neuropil in the fly brain. Representative images show the central neuropil in paraffin-embedded, HE-stained brain sections from 1-day-old or 7-week-old flies. Scale bars: 500 μm. The percentages of the vacuolar areas (indicated by the arrows in the images) in the central neuropil are shown. n = 8-12 hemispheres; ***p < 0.001 by Student’s *t*-test.

In *Drosophila*, brain vacuolation is a morphological hallmark of neurodegeneration. We next examined structural changes in the fly brains using serial sections of whole brains. In aged (49-day-old) fly brains expressing control RNAi targeting mCherry, neurodegeneration was spontaneously observed (**Figure 5D**). However, neuronal knockdown of *Rac1* induced prominent neurodegeneration in the neuropil region of aged (49-day-old) but not young (1-day-old) fly brains (**Figure 5D**). These results suggest that neuronal knockdown of *Rac1* is sufficient to cause age-dependent neurodegeneration in flies.

To minimize potential off-target effects of RNAi, histological analysis was performed in an independent transgenic line carrying RNAi targeting a different region of *Rac1* (**Figure S4**, TRiP RNAi line #28985). Although the knockdown efficiency of *Rac1* by this RNAi was decreased (**Figure S4A**), age-dependent neurodegeneration was similarly observed in these fly brains (**Figure S4B**).

### miRNA hsa-miR-101-3p represses *RAC1* expression

To gain insight into the potential mechanism underlying the downregulation of *RAC1* expression in the EC, we focused on miRNAs, which are short non-coding RNA molecules of 21-25 nucleotides that bind to complementary mRNAs to inhibit mRNA translation. We searched for candidate miRNAs that inhibit *RAC1* expression using starBase, a database for searching candidate miRNA-mRNA interactions from experimental data sets (crosslinking immunoprecipitation (CLIP)-seq or degradome-seq) and seven prediction tools. More than three prediction tools identified that 62 miRNAs could bind to *RAC1.* Furthermore, more than five experimental data sets supported the interactions between the associated miRNAs and *RAC1*.

To narrow down candidate miRNAs, we measured comprehensive miRNA expression levels using small RNA sequencing in brain tissues obtained from the same individuals used for microarray analysis (40 samples in each brain region). We found nine miRNAs whose expression levels were negatively correlated with *RAC1* gene expression in the EC (Spearman correlation coefficient (SCC)<0; **Table 1**). Among these miRNAs, hsa-miR-101-3p showed the strongest negative correlation (SCC=-0.542), and its interaction with *RAC1* was validated in the most experimental data sets (**Figure 6A**). The SCCs between *RAC1* and hsa-miR-101-3p also indicated a negative correlation in the TC and FC (SCC = −0.644 in the TC; SCC = −0.336 in the FC). Moreover, the hsa-miR-101-3p levels were significantly increased with progressive Braak NFT stage in the EC (**Figure 6B**, SCC = 0.491; p-value = 1.29×10^−3^), suggesting that this miRNA responded to disease progression.

**Figure 6.**
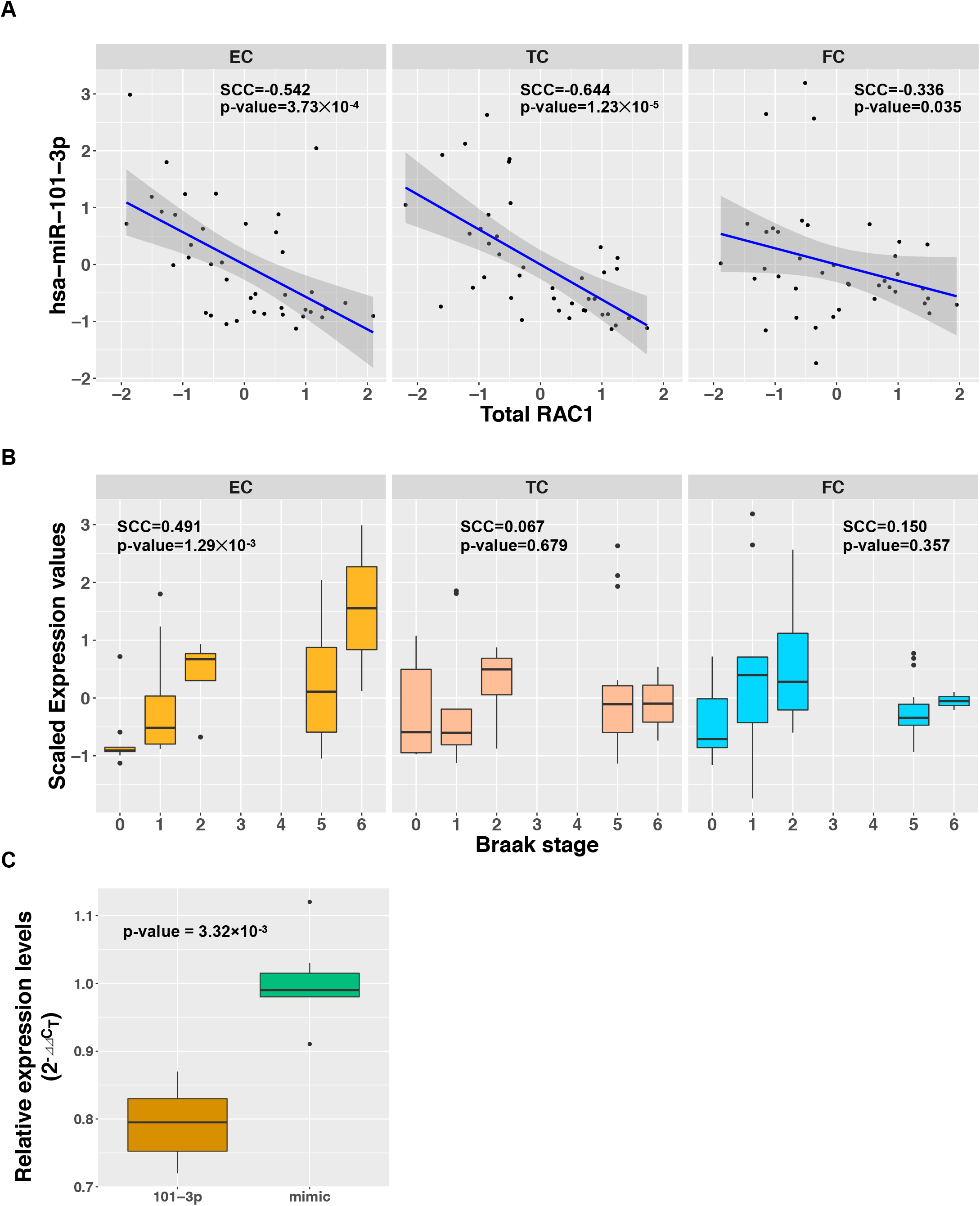
hsa-miR-101-3p represses *RAC1* expression. (A) Correlation of hsa-miR-101-3p expression across Braak stages. The p-values were calculated by a test for no correlation. (B) Correlation between the hsa-miR-101-3p and total RAC1 levels in each brain region. (C) *RAC1* expression levels in SH-SY5Y neuronal cells overexpressing hsa-miR-101-3p. *RAC1* expression levels were quantified by qRT-PCR and were normalized to *GUSB* expression. The p-value was calculated by the Wilcoxon rank sum test.

**Table1.** Candidates of miRNAs targeting RAC1.

| Table1. Candidates of miRNAs targeting RAC1. |  |  |  |  |  |  |  |
| --- | --- | --- | --- | --- | --- | --- | --- |
| miRNAname | Binding region | #CLIP-Seq | #Degradome-Seq | #Database Hits | RAC1-miRNA expression correlation |  |  |
|  |  |  |  |  | EC | TC | FC |
| hsa-miR-101-3p | chr7:6442614-6442620 [+] | 36 | 1 | 4 | -0.542 | -0.644 | -0.336 |
| hsa-miR-365a-3p | chr7:6442981-6442987 [+] | 30 | 0 | 5 | -0.286 | -0.108 | 0.037 |
| hsa-miR-365b-3p | chr7:6442981-6442987 [+] | 30 | 0 | 3 | -0.286 | -0.108 | 0.037 |
| hsa-miR-155-5p | chr7:6443104-6443109 [+] | 19 | 0 | 4 | -0.390 | -0.027 | 0.085 |
| hsa-miR-155-5p | chr7:6443554-6443560 [+] | 17 | 0 | 4 | -0.390 | -0.027 | 0.085 |
| hsa-miR-194-5p | chr7:6443146-6443151 [+] | 16 | 0 | 3 | -0.412 | -0.394 | -0.164 |
| hsa-miR-194-5p | chr7:6442204-6442209 [+] | 12 | 0 | 5 | -0.412 | -0.394 | -0.164 |
| hsa-miR-495-3p | chr7:6442202-6442207 [+] | 12 | 0 | 3 | -0.302 | -0.182 | -0.147 |
| hsa-miR-22-3p | chr7:6443216-6443221 [+] | 11 | 0 | 3 | -0.370 | -0.503 | -0.066 |
| hsa-miR-1296-5p | chr7:6442312-6442317 [+] | 8 | 1 | 3 | -0.311 | 0.084 | -0.120 |
| hsa-miR-495-3p | chr7:6442264-6442268 [+] | 5 | 0 | 4 | -0.302 | -0.182 | -0.147 |
| "#CLIP-Seq" and "#Degradome-Seq" indicate the number of experiments that found miRNA-RAC1 interactions using CLIP-Seq and Degradome-Seq, respectively. "#Database Hits" indicates the number of databases that predicted miRNA-RAC1 interactions in seven databases (PITA, RNA22, miRmap, microT, miRanda, PicTar, and TargetScan). "RAC1-miRNA expression correlation" shows Spearman correlation coefficients between RAC1 expression and miRNA expression in our data. miRNAs with #CLIP-Seq ≥ 5, #Database Hits ≥ 3, and Spearman correlation coefficient in EC < 0 were selected. |  |  |  |  |  |  |  |

To experimentally validate the suppression of *RAC1* expression by hsa-miR-101-3p, we transfected human neuronal SH-SY5Y cells with hsa-miR-101-3p and quantified *RAC1* expression by quantitative reverse transcription PCR (qRT-PCR). Overexpression of hsa-miR-101-3p resulted in a significant decrease in the *RAC1* gene expression level compared to that in mimic-transfected control cells (p-value = 3.32×10^−3^), indicating that hsa-miR-101-3p regulates *RAC1* in human neuronal cells (**Figure 6C**). Taken together, these results suggest that the increased expression of hsa-miR-101-3p with progressive Braak NFT stage may be involved in the suppression of *RAC1* expression in AD brains.

## Discussion

Several network analyses detected alterations in biological networks in AD brains (10,11,35,36). In this study, to analyse the dynamics of molecular networks during the progression of Braak NFT stage, we constructed PDNs in three regions of AD brains at each Braak NFT stage (**Figures 2 and 3**). Our analyses revealed that both gain and loss of PDNs occurred with the progression of Braak NFT stage (**Figure 3**) and that loss of PDNs was more prominent than gain of PDNs in all three brain regions. These data are consistent with our previous findings (35) and provide further evidence that PDNs collapse during disease progression, which may underlie the neuronal dysfunction and neurodegeneration in AD.

Consistent with the severity of brain pathology (1), our analysis detected that the EC was the most severely affected among the three brain regions analysed (**Figure 3C**). Furthermore, gene functional enrichment analysis revealed that genes in appearing or disappearing PDNs in the EC were associated with functions related to immune systems and phagocytosis. Alterations in these pathways have been implicated in brain pathologies in AD (37–42). Moreover, recent GWASs in AD identified the SNPs located in the coding regions of or in the vicinity of genes associated with immune functions, suggesting that alterations in the immune system play significant roles in AD pathogenesis (43–46). We identified Integrin Subunit Alpha X (*ITGAX*), also known as *CD11c*, as an immune gene in appearing PDNs in the EC (**Table S2**). *ITGAX/CD11c* expression is elevated in the EC and the hippocampus in AD (47), and recent studies found that *ITGAX/CD11c* is a marker of disease-associated microglia (DAM) in late-stage neurodegenerative diseases, including AD (48,49). Indeed, *ITGAX/CD11c* was expressed only in Braak stages V+VI in our data sets, suggesting that the formation of DAM is activated in the EC in the late stage of AD.

Our results strongly suggest that loss of PDNs is more prominent than gain of PDNs during the progression of NFT pathology (**Figure 3**). Among these 15 collapsed genes, we identified 5 genes, including *RAC1*, as hub genes, which play key roles in the alteration of the PDNs. Using independent data sets, we confirmed that the mRNA expression level of *RAC1* was downregulated in the EC of AD brains (**Figure 4**). Importantly, these changes are not simply due to ageing effects. A previous study showed that *RAC1* expression was negatively correlated with phosphorylated tau in the frontal lobes of AD patients (50). In contrast, amyloid-β (Aβ) pathology was not correlated with *RAC1* expression (50), and we found no correlation between *RAC1* expression and Braak SP stage (FDR-adjusted p-value=0.337). Interestingly, *RAC1* plays a key role in the phagocytosis process, and its dysfunction impairs the removal of synapses, cellular debris and aggregated proteins (51,52). These reports suggest that the collapse of a *RAC1*-centred PDN may promote synaptic dysfunction and neurodegeneration in AD.

To examine the effects of *RAC1* downregulation on neural integrity during ageing, we utilized the fruit fly *Drosophila* as an *in vivo* genetic model. *RAC1* activity is involved in the formation and maintenance of memory in flies (53) and mice (54–56). Moreover, *RAC1* dysfunction is implicated in several neurodegenerative diseases, such as Parkinson’s disease and amyotrophic lateral sclerosis (57–60). We found that neuronal knockdown of the *Drosophila* orthologue of human *RAC1, Rac1*, was sufficient to induce age-dependent behavioural deficits and neurodegeneration (**Figure 5**). These results suggest that our analysis identified one of the neurodegenerative processes associated with NFT pathology in AD pathogenesis.

Finally, we identified a miRNA, hsa-miR-101-3p, as a potential regulator of *RAC1* in AD brains (**Figure 6**). As Braak NFT stage progressed, the expression of hsa-miR-101-3p was upregulated specifically in the EC. Furthermore, overexpression of hsa-miR-101-3p in the human neuronal cell line SH-SY5Y caused *RAC1* downregulation. Interestingly, recent studies have reported that hsa-miR-101-3p is differentially expressed in AD serum (61), and precursor as well as mature forms of miR-101 regulate the expression of amyloid-β precursor protein in human HeLa cells and rat hippocampal neurons (62,63). miR-101 is also associated with the innate immune response in macrophages (64) and hepatocellular carcinoma (65). These reports further suggest the potential involvement of has-miR-101-3p in AD pathogenesis.

In conclusion, we demonstrated that disruption of a *RAC1*-centred PDN is associated with NFT pathology in AD brains and is sufficient to cause age-dependent neurodegeneration. This study suggests the utility of our integrated network approach for revealing the mechanisms underlying AD and identifying potential therapeutic targets.

## Materials and Methods

### Postmortem brain tissues

All 71 Japanese subjects were recruited at Niigata University. These subjects were included in a previous study (7). Based on the Braak NFT staging system established by Braak H and Braak E (1), the subjects were divided into four groups: Braak NFT stage 0 (N=13), I-II (N=20), III-IV (N = 19) and V-VI (N = 19). Three brain regions (the EC, TC and FC) were used. In total, 213 brain tissue specimens (= 71 subjects×3 brain regions) were ultimately included in this study. Detailed statistical information about the samples is described in our previous study (7).

### Total RNA extraction and quality control

Total RNA was extracted from brain tissues and cultured cells with a TRIzol Plus RNA Purification System (Life Technologies, Carlsbad, CA, USA). Genomic DNA was removed through on-column DNase I treatment during RNA preparation. To determine the RNA integrity number (1 (totally degraded) to 10 (intact)), 11 a 2100 Bioanalyzer instrument was used with an RNA 6000 Pico Assay (Agilent, Santa Clara, CA, USA). We fluorometrically determined the concentration of total RNA with a Quant-iT RiboGreen RNA Assay Kit (Life Technologies).

### Exon-level gene expression profiling

We used GeneChip Human Exon 1.0 ST arrays (Affymetrix, Santa Clara, CA, USA) to measure the expression levels of transcripts in each brain tissue specimen. The initial raw data (DAT files) were processed into CEL files via Affymetrix GeneChip Operating Software. Partek Genomics Suite 6.4 (Partek, St. Louis, MO, USA) was used to normalize the data in the CEL files and summarize the expression of the probesets in the exon array. Background correction was performed via a robust multi-array average (RMA) method (66) with adjustment for GC content. The expression levels of all probesets in the CEL files were log2-transformed. A total of 878,018 probesets were used. To filter out low expression signals (including noise or poorly hybridized probes), which may lead to false positives, the DABG p-values of the exon probesets were calculated using Affymetrix Power Tools (APT, http://www.affymetrix.com/partners_programs/programs/developer/tools/powertools.affx). The probesets and transcripts were annotated according to the human genome hg19 reference sequence.

### Domain-domain interaction data set

We used domain-domain interactions registered in INstruct (17). This data set includes 3,626 proteins and 11,470 domain-domain interactions in *Homo sapiens.* We used INstruct-treated protein interactions that were validated through eight protein interaction databases (BioGrid (67), Database of Interacting Proteins (DIP) (68), Human Protein Reference Database (HPRD) (69), IntAct (70), iRefWeb (71), The Molecular Interaction Database (MINT) (72), Munich Information Center for Protein Sequences (MIPS) (73) and VisAnt (74)) and curated domain-domain interactions that were supported by co-crystal structures. We removed 3,414 self-interactions.

### Estimation of domain expression levels

We indirectly inferred the expression levels of protein domains using the expression levels of transcripts measured in the exon array. The probes in the exon array were assigned to the exons of each gene based on the positions of each exon using refGene.txt provided by the UCSC Genome Browser. We defined an expressed exon as an exon with a median DABG p-value of < 0.05 and a median log2-transformed expression value of ≥6 across all samples in a Braak NFT stage in a brain region. To determine the exons constructing the domains, we linked the coding sequences (CDSs) of the cDNAs and amino acid sequences of the proteins. The sequence information of the CDSs was obtained from the RefSeq database (human.rna.gbff). For amino acid sequences, information from the UniProt database (uniprot_sprot.fasta) was used, because the protein domains in INstruct were based on the canonical protein sequences in the UniProt database. The length of a CDS should be three times that of the corresponding amino acid sequence. We confirmed this criterion for all genes/proteins targeted in this study using the above data. Using this correspondence between the CDSs and the protein sequences, we searched the exons that were included in the domains. If an exon region covered more than half of a domain region or a domain region covered more than half of an exon region, we considered the exon to be a component of the domain.

### Calculation of theoretical values

The theoretical value *T* for the number of interactions in an expression pattern was calculated based on the probabilities of interactions expressed in each Braak NFT stage:

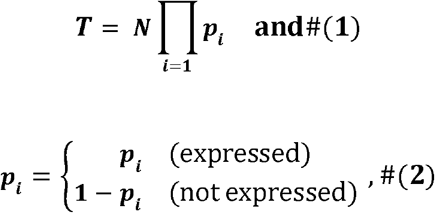

where *N* is the total number of interactions expressed in at least one Braak NFT stage in a brain region, and *p_i_* indicates an observed probability of an interaction expressed in Braak NFT stage *i*. The observed probability was calculated as the number of interactions expressed in Braak NFT stage *i* divided by *N*. For instance, the theoretical value of “Pattern 1” in disappearing interactions shown in **Figure 3A** was calculated as 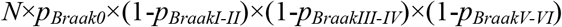.

### Generation of null distribution

To test the statistical significance of the number of interactions in each expression pattern, we generated null distributions corresponding to each expression pattern shown in **Figure 3A**. The null distributions were generated by “expression label shuffling” in each Braak NFT stage (**Supplementary Figure 3**). This procedure involved a set of shuffles from Braak stage 0 to Braak stages V-VI as one trial and was repeated 10,000 times. We calculated the numbers of interactions in each expression pattern in each trial and generated null distributions for each expression pattern. To assess the statistical significance of each expression pattern, we calculated the z-score as *Z*=(*x-μ*)/*σ*, where *x* is the observed number of interactions in an expression pattern and *μ* and *σ* are the mean value and standard deviation calculated from the null distribution of the expression pattern, respectively. The values of *μ* converge to the theoretical values described above. Each z-score calculated for each expression pattern was converted to a p-value.

### Gene functional enrichment analysis

We used Metascape to examine the functions of genes in appearing or disappearing PDNs (75). Terms with a q-value of < 0.01, a gene count of > 3, and an enrichment factor of > 1.5 (the enrichment factor is the ratio of the observed count to the count expected by chance) were assembled and grouped into clusters based on their membership similarities.

### Estimation of the effects of Braak NFT stage on domain expression

To examine whether the expression levels of domains depend on Braak NFT stage rather than ageing, we used the following linear regression model:

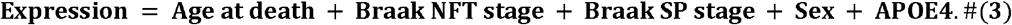

Here, Braak SP stage is represented as 0, A, B, or C. These stages were converted to countable values (0∼3). Sex was treated as a categorical variable. APOE4 represents the number of Apoe ε4 alleles (0, 1, or 2). These variables were analysed by a Z-test using the R statistical package glm.

### Validation of *RAC1* expression changes using independent data sets

We used AlzData to obtain gene expression levels in publicly available data sets (76). AlzData is a database storing statistical meta-analysis results using 78 samples from four data sets in the EC. Gene expression profiles were analysed by a linear regression model with age and sex as covariates. Log2 fold change (log2FC) values and FDR-adjusted p-values were calculated in AlzData. We downloaded the expression profiles from AlzData and calculated correlation coefficients between the expression levels of the genes and the age of the samples.

### *Drosophila* genetics

Flies were maintained in standard cornmeal media at 25 °C. Elav-GAL4 (#458), UAS-*mcherry* RNAi (#35785) and UAS-*Rac1* RNAi (#34910 and #28985) transgenic flies were obtained from the Bloomington Stock Center. Experiments were performed using age-matched male flies. The ages of the flies used in the experiments presented in Figure 5A and Figure S4A were 7 days old, and the ages of the other flies are indicated in the figures. The genotypes of the flies were as follows: (*mcherry* RNAi): elav-GAL4/Y;;UAS-*mcherry* RNAi/+, (*Rac1* RNAi): elav-GAL4/Y;;UAS-*Rac1* RNAi #34910/+, and (*Rac1* RNAi (#28985)): elav-GAL4/Y;;UAS-*Rac1* RNAi #28985/+.

### RNA extraction and quantitative real-time PCR analysis for *Drosophila*

More than 25 flies of each genotype were collected and frozen. The heads were mechanically isolated, and total RNA was extracted using TRIzol reagent (Thermo Fisher Scientific) according to the manufacturer’s protocol, with an additional centrifugation step (16,000 × g for 10 min) to remove cuticle membranes prior to the addition of chloroform. Total RNA was reverse transcribed using a PrimeScript RT-PCR kit (TaKaRa Bio), and qRT-PCR was performed using SsoAdvanced Universal SYBR Green Supermix (Bio-Rad Laboratories) on a CFX96 real-time PCR detection system (Bio-Rad Laboratories). The average threshold cycle value was calculated from at least three replicates per sample. The expression of genes of interest was normalized to GAPDH1 expression. The results are presented as the means ± s.e.m. An unpaired Student’s *t*-test (Prism 7, GraphPad) was used to determine statistical significance. The primer sequences used in this study are shown below:

*GAPDH1* (*Drosophila*), 109 bp Forward; 5’-GACGAAATCAAGGCTAAGGTCG-3’ Reverse; 5’-AATGGGTGTCGCTGAAGAAGTC-3’
*Rac1* (*Drosophila*), 199 bp Forward; 5’-ACGTTCCATTGCAATTACACTTAC-3’ Reverse; 5’-AGTTGCTCAGTTTGATGCCT-3’

### Climbing assay

The climbing assay was performed as described previously (33,34). Approximately 25 male flies were placed in an empty plastic vial. The vial was then gently tapped to knock all flies to the bottom. The flies in the top, middle, and bottom third of the vial were counted after 10 seconds. The percentage of flies that stayed at the bottom was used for statistical analyses (unpaired Student’s *t*-test).

### Life span analysis

Life span analysis was performed as described previously (33,34). Food vials containing 25 flies were placed on their sides at 25 °C under conditions of 70% humidity and a 12:12-h light:dark cycle. Food vials were changed every 3-4 days, and the number of dead flies was counted at each change. At least four vials per genotype were prepared. Experiments were repeated more than three times, and a representative result is shown. Kaplan-Meier survival analysis with a log-rank test (SigmaPlot 11.0, Systat Software, Inc.) was used to determine statistical significance for life span analysis data.

### Histological analysis

The heads of male flies were fixed in 4% paraformaldehyde for 24 h at 4 °C and embedded in paraffin. Serial sections (6 μm thickness) through the entire head were prepared, stained with haematoxylin and eosin (Sigma-Aldrich), and examined by bright field microscopy. Images of the sections were acquired with an AxioCam 105 colour camera (Carl Zeiss). For quantification, we focused on analysing sections covering central neuropil regions. We selected the section with the most severe neurodegeneration in each individual fly, and the vacuolar area was measured using ImageJ (NIH). The results are expressed as the means ± s.e.m. An unpaired Student’s *t*-test was used to determine statistical significance.

### Searching of candidate miRNAs targeting *RAC1* mRNA

We used starBase to find candidate miRNAs that target *RAC1* mRNA (77). StarBase is a database for searching candidate miRNA-mRNA interactions from experimental data sets (CLIP-seq or degradome-seq) and seven prediction databases (Probability of Interaction by Target Accessibility (PITA) (78), RNA22 (79), miRmap (80), microT (81), miRanda (82), PicTar (83), and TargetScan (84)). We selected the miRNAs with validation by CLIP-Seq in ≥ 5 studies, ≥ 3 database hits, and a Spearman correlation coefficient in the EC of < 0.

### miRNA overexpression in cultured cells

The human neuroblastoma cell line SH-SY5Y was used for miRNA overexpression experiments. Cell culture and miRNA transfection were performed as described previously (85). SH-SY5Y cells were transfected with either the hsa-miR-101-3p mimic (MC11414; Thermo Fisher Scientific, Waltham, MA, USA) or scrambled control (mirVana™ miRNA Mimic, Negative Control #1; Thermo Fisher Scientific). Cells were harvested using TRIzol reagent 24 h after transfection and stored at −80 °C until RNA extraction. A total of 200 ng of total RNA was reverse transcribed to cDNA using SuperScript IV VILO Master Mix (Thermo Fisher Scientific) in a reaction volume of 20 μl. The cDNA product was diluted by adding 60 μl of water. qPCR was performed in triplicate, with 2.5 μl of the diluted product in a reaction volume of 10 μl in a 384-well plate on an ABI PRISM 7900HT (Thermo Fisher Scientific) instrument with TaqMan Gene Expression Assays (Hs01902432_s1 for *RAC1* and Hs99999908_m1 for *GUSB*; Thermo Fisher Scientific). The relative gene expression levels of *RAC1* were calculated by the 2^−ΔΔCT^ method using *GUSB* as the endogenous control for normalization. Six biological replicates were included for each condition, and the statistical significance between the two conditions was analysed by the Wilcoxon rank sum test.

## Accession number

Gene expression data are available in the GEO database (GEO accession number GSE131617).

## Acknowledgements

Certain pictures in the figures were obtained from the Togo picture gallery (https://togotv.dbcls.jp/pics.html). We acknowledge all groups that have contributed.

## Abbreviations

AD: Alzheimer’s disease
NFT: neurofibrillary tangle
EC: entorhinal cortex
TC: temporal cortex
FC: frontal cortex
PDN: protein domain network

---

**Figure S1.**
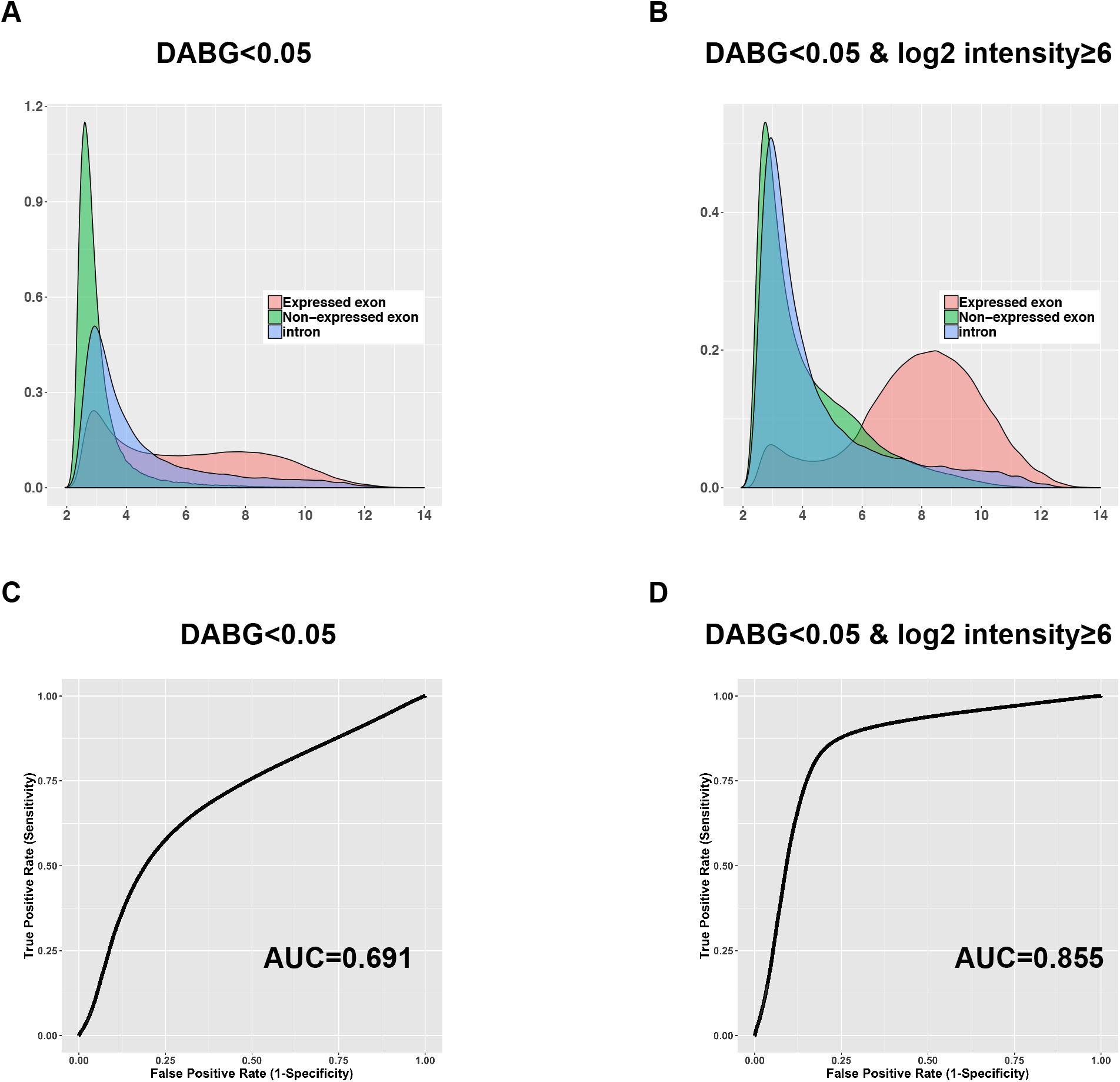
Discrimination between the expressed exons and the intron. Density distributions of when using only DABG P-value (A) and when using DABG P-value and a cutoff value of 6 (B). ROC curves of when using only DABG P-value (C) and when using DABG P-value and a cutoff value of 6 (D).

**Figure S2.**
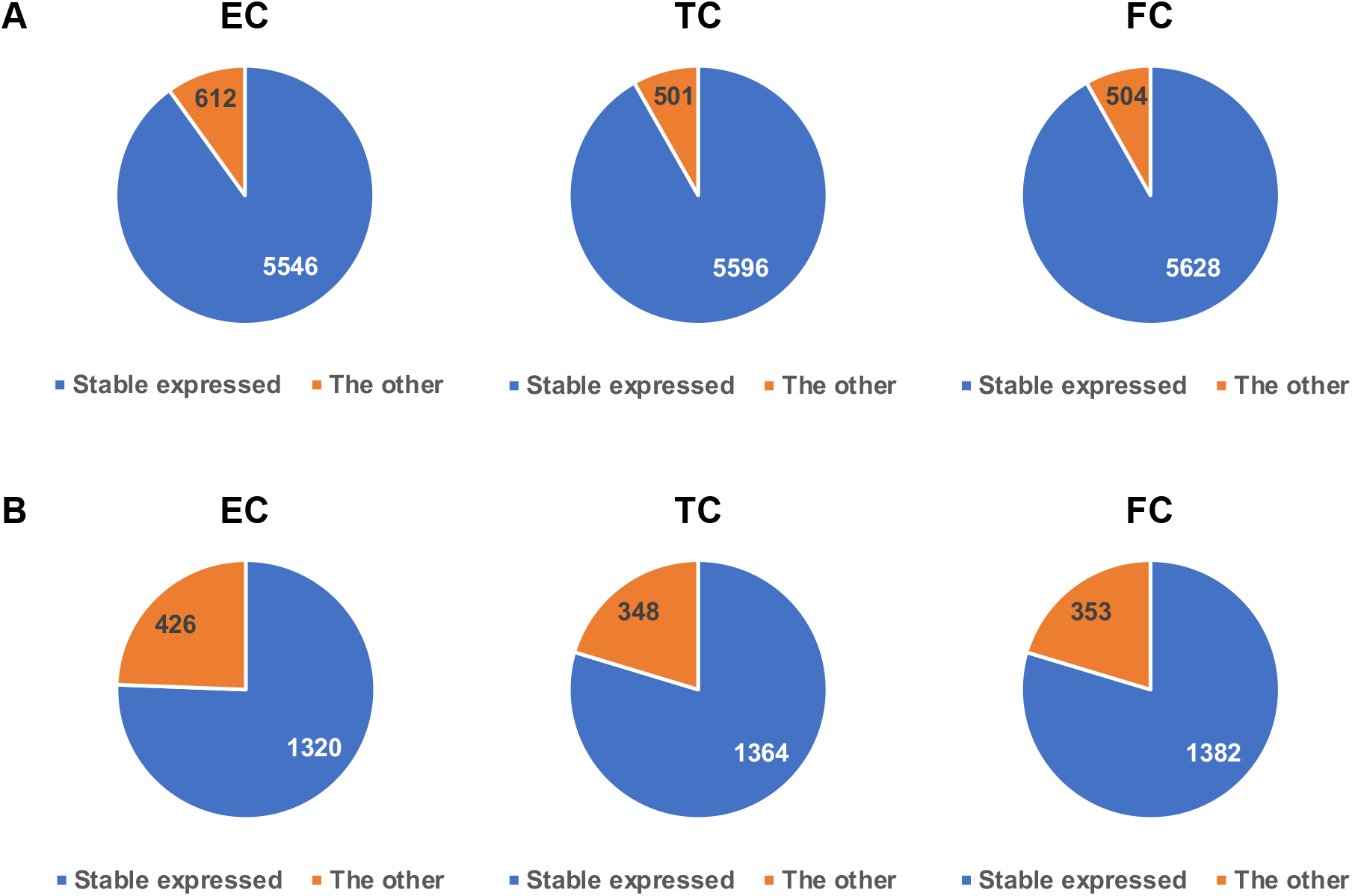
The influence of exon usages. Pie charts of the expressed exons (A) and the proteins with one or more unstable expressed domains (B).

**Figure S3.**
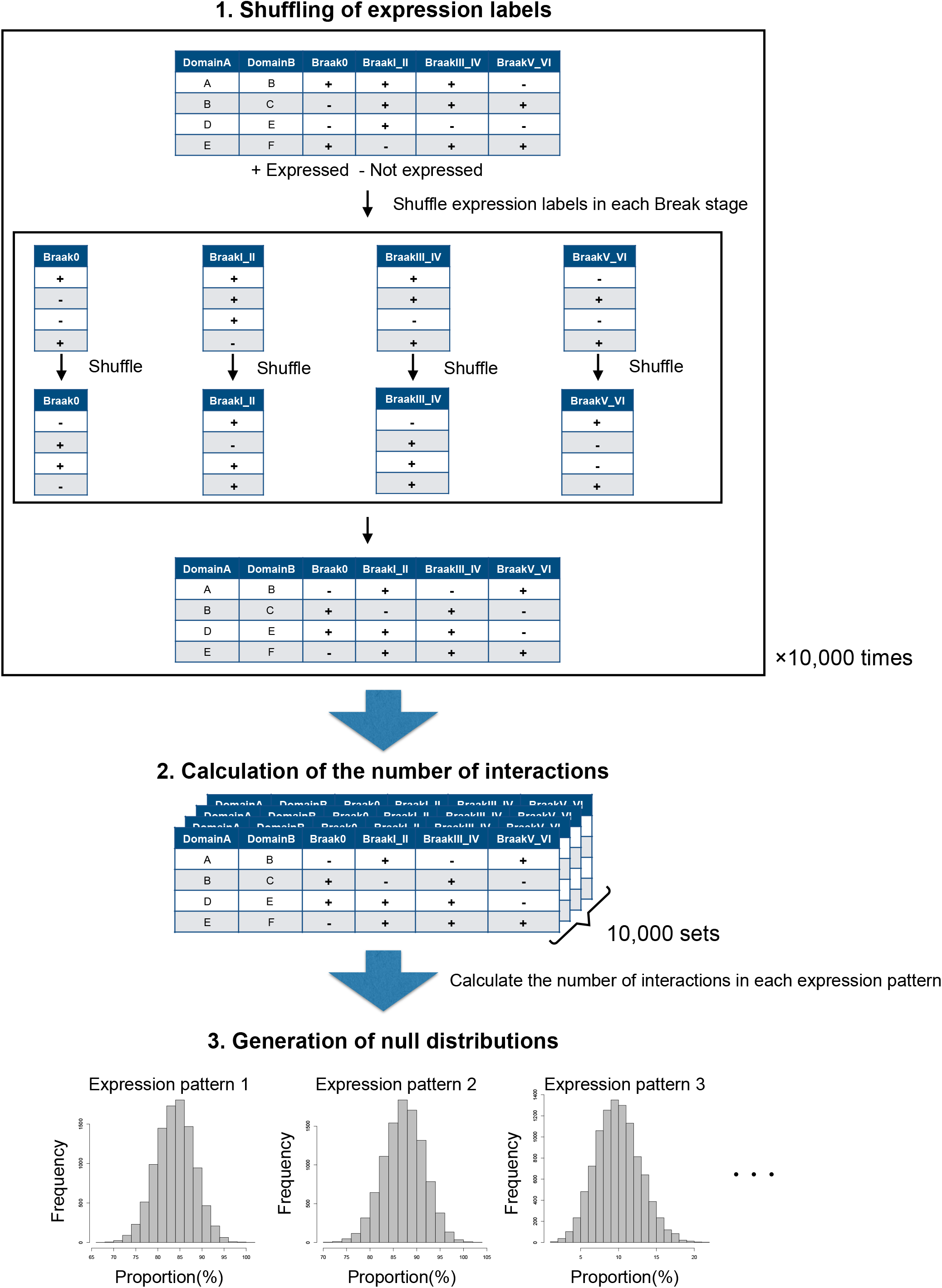
Generation of null distributions in each expression pattern. We performed three steps to obtain null distributions. In the first step, we shuffled expression labels in each Break stage. After shuffling, we merged them in each interaction. These procedures were repeated 10,000 times. In the second step, we collected the 10,000 shuffled sets and calculated the proportion of interactions in each expression pattern. Finally, we obtained the null distributions in each expression pattern.

**Figure S4.**
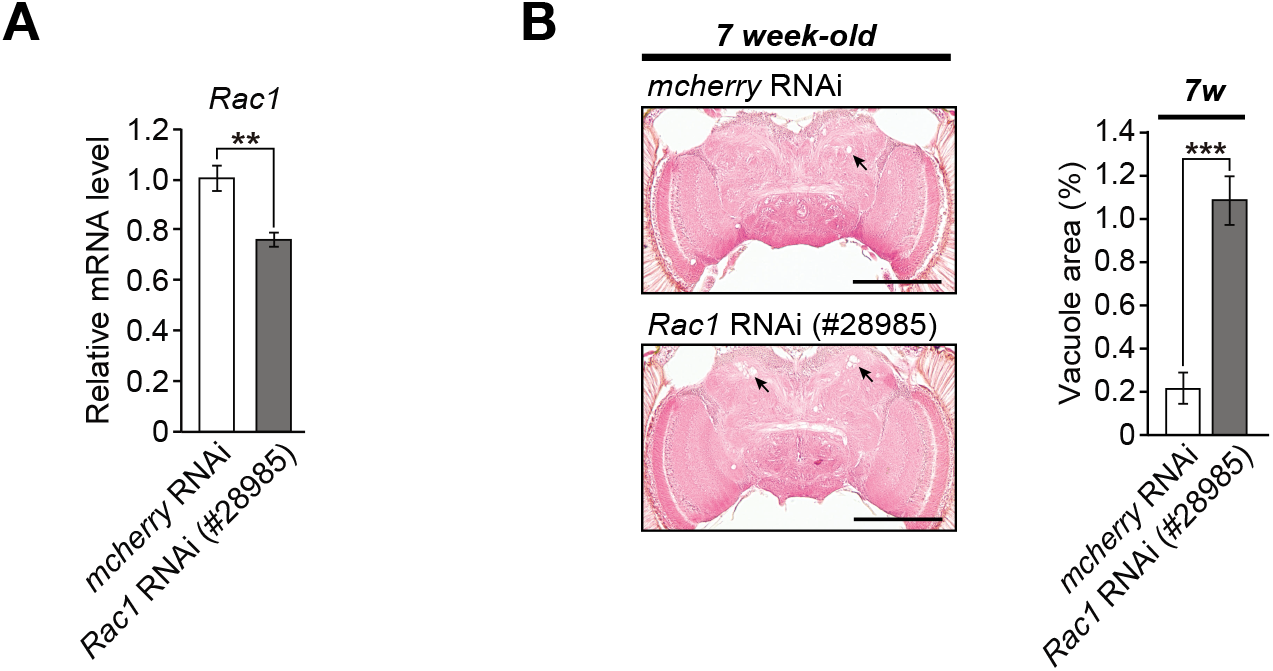
Neuronal knockdown of *Rac1* (#28985) causes age dependent neurodegeneration. (A) mRNA levels of *Rac1* (#28985) in heads of flies carrying the RNAi transgene targeting *Rac1* were analysed by qRT-PCR. n = 4, **p < 0.01 by Student’s *t*-test. (B) Knockdown of *Rac1* (#28985) in neurons caused age-dependent neurodegeneration in the calyx (dendrite) region of central neuropil in the fly brain. Representative images show the central neuropil in paraffin-embedded, HE-stained brain sections from 7-week-old flies. Scale bars: 500 μm. The percentages of the vacuolar areas (indicated by the arrows in the images) in central neuropil are shown. n = 12 hemispheres; ***p < 0.001 by Student’s *t*-test. The genotypes of the flies were as follows: (*mcherry* RNAi): elav-GAL4/Y;; UAS-*mcherry* RNAi/+, and (*Rac1* RNAi (#28985)): elav-GAL4/Y;; UAS-*Rac1* RNAi/+. The age of the flies used in (A) were 7-day-old.

